# Comprehensive Classification of the RNase H-like domain-containing Proteins in Plants

**DOI:** 10.1101/572842

**Authors:** Shuai Li, Kunpeng Liu, Qianwen Sun

## Abstract

**Background:** R-loop is a nucleic acid structure containing an RNA-DNA hybrid and a displaced single-stranded DNA. Recently, accumulated evidence showed that R-loops widely present in various organisms’ genomes and are involved in many physiological processes, including DNA replication, RNA transcription, and DNA repair. RNase H-like superfamily (RNHLS) domain-containing proteins, such as RNase H enzymes, are essential in restricting R-loop levels. However, little is known about the function and relationship of other RNHLS proteins on R-loop regulation, especially in plants.

**Results:** In this study, we characterized 6193 RNHLS proteins from 13 representative plant species and clustered these proteins into 27 clusters, among which reverse transcriptases and exonucleases are the two largest groups. Moreover, we found 691 RNHLS proteins in Arabidopsis with a conserved catalytic alpha-helix and beta-sheet motif. Interestingly, each of the Arabidopsis RNHLS proteins is composed of not only an RNHLS domain but also another different protein domain. Additionally, the *RNHLS* genes are highly expressed in different meristems and metabolic tissues, which indicate that the RNHLS proteins might play important roles in the development and maintenance of these tissues.

**Conclusions:** In summary, we systematically analyzed RNHL proteins in plants and found that there are mainly 27 subclusters of them. Most of these proteins might be implicated in DNA replication, RNA transcription, and nucleic acid degradation. We classified and characterized the RNHLS proteins in plants, which may afford new insights into the investigation of novel regulatory mechanisms and functions of R-loops.

## Introduction

R-loop is a nucleotide structure that consists of an RNA-DNA hybrid and a displaced single-stranded DNA [1]. Previously, R-loops have been considered as transcription by-products since nascent transcripts could bind back to their DNA templates, which could lead to genome instability [2]. Nonetheless, recent studies in different organisms revealed that R-loops are prevalent and widespread on genomes of yeasts, plants, and mammals [3–9]. Besides, R-loops are involved in DNA replication, retrovirus infection, and retrotransposon life cycle *in vivo* [10]. These hints together suggested that R-loops should be accurately modulated in different cellular processes and tissues.

Ribonucleases Hs (RNase H), a type of endonuclease, can remove R-loops through degrading the RNA moiety of an RNA-DNA hybrid [11–13]. It was suggested that RNA molecules might be the origin of life, while RNase H enzymes mainly disrupt direct RNA-DNA interactions [14]. Thus, it is reasonable that RNase H might be one of the most ancient proteins and many enzymes involved in nucleotide-related processes might employ the RNase H domain during evolution [15, 16]. RNase H-like superfamily (RNHLS) proteins are a group of proteins with similar 3D structure of RNase H domain, including type 1 and type 2 RNase Hs, retroviral integrases, DNA transposases, Holliday junction resolvases, Piwi/Argonaute nucleases, exonucleases, key spliceosome component Pre-mRNA-processing-splicing factor 8 (Prp8), and other RNA nucleases [17–23]. Decades of research have revealed that RNHLS proteins play critical roles in diverse cellular processes, such as DNA replication and repair, RNA transcription and interference, homologous and non-homologous recombination, transposon and retrovirus transposition, and single-stranded RNA virus infection [15]. A previous study using bioinformatic approaches to search protein databases for all RNHLS proteins available revealed that RNHLS proteins could be grouped into 152 families and principally divided into exonucleases and endonucleases [15].

Plants are sessile and thus need to employ different strategies, such as responses induced by various hormones and metabolites, to survive at different biotic or abiotic stresses. R-loops might play an important role in these responses. Recently, it was reported that R-loop levels could impact an antisense transcription at *FLC* locus in Arabidopsis, which represses the accumulation of the antisense long non-coding RNA [24]. Moreover, the misexpression of topoisomerase 1, a critical R-loop regulator, could compromise the expression of auxin-related genes in rice [25]. In addition, there are three RNase H1 enzymes with different organelle localization and the chloroplast-localized RNase H1 is critical for maintaining chloroplast genome integrity in Arabidopsis [26]. Furthermore, R-loops can perform their function as chromatin features and read by proteins such as ALBA proteins in Arabidopsis and GADD45A (growth arrest and DNA damage inducible alpha) proteins in mammals [27, 28]. Therefore, RNHLS proteins might regulate R-loops in development tissues and response to hormones in plants whose gene expression should be fine-tuned. However, little is known about RNHLS proteins and their relationship in plants, especially on the regulation of R-loops.

In this study, we systemically analyzed the relationship and diversity of RNHLS proteins *in silico* in plants. We searched databases and collected 6193 RNHLS proteins from 13 representative plant species, including *Solanum lycopersicum* (tomato), *Solanum tuberosum* (potato), *Populus trichocarpa* (western balsam poplar), *Glycine max* (soybean), *Citrus sinensis* (sweet orange), *Arabidopsis thaliana* (arabidopsis), *Oryza sativa* (rice), *Physcomitrella patens* (moss), *Marchantia polymorpha* (liverwort), *Klebsormidium flaccidum* (filamentous charophyte green algae), *Chlamydomonas eustigma* (unicellular flagellate green algae), *Cyanidioschyzon merolae* (red alga), and *Chondrus crispus* (carrageen Irish moss). Moreover, these proteins could be clustered into 27 clusters, of which reverse transcriptases and exonucleases are the two largest groups. Besides, we identified 691 RNHLS proteins in Arabidopsis, most of which are combined with different protein domains, and found that the catalytic alpha-helix and the beta-sheet motifs are conserved among them. Additionally, we showed that the *RNHLS* genes not only are actively expressed in meristems and other metabolic tissues but also could be grouped together. In summary, these results suggested that the RNHLS proteins play essential roles in plants, especially on the development and maintenance of meristems and metabolic tissues.

## Materials and methods

### Co-occurrence of RNHLS domain with other protein domains

The co-occurrence matrix of the RNHLS domain (3.30.420.10) with other protein domains was downloaded from Gene3D [29]. The heatmap was processed by online software Morpheus (https://software.broadinstitute.org/morpheus/) using the default settings.

### Protein sequence search

To identify the RNase H-like superfamily proteins in Arabidopsis comprehensively, we used the proteins sequences from RNHLS (IPR012337) of the InterPro database (http://www.ebi.ac.uk/interpro/) filtered with different species. Also, we performed sequence BLAST with AtRNH1A (At3g01410) as a query in the Phytozome and NCBI database [30, 31]. We carried out the analysis after combining the sequence from the three methods and removed duplicates.

### Protein sequence cluster

Identified proteins were first analyzed using CD-HIT with a threshold at 60 percent and word size of 3 [32]. The clustered proteins were aligned using the BLAST to compute and evaluate the similarity with an average pairwise E-value at least 1×10^−18^ [31]. The network was visualized using Cytoscape3.7.0 and laid out using the organic layout algorithm and annotated manually [33].

### Protein sequence alignment

To analysis the protein sequences systemically, we used Clustal Omega online service with the default parameters except for all iterations at five [34]. Alignment results were visualized with UGENE and MEGA7 [35]. Structure-Based sequence alignment was performed to find the conserved secondary structure using the PROMALS3D and JPred4 and then manually adjusted with UGENE with shifting insertions and deletions to identify the homologous positions from conserved secondary structure elements [36, 37].

### Phylogenetic tree construction

The phylogenetic tree of the subgroups and Arabidopsis RNHLS proteins were generated using Clustal Omega at the same parameters for protein sequence alignment. For the phylogenetic tree of the total RNHLS proteins, we employed the MAFFT for multiple sequence alignment and constructed the phylogenetic tree using FastTree [38, 39]. We further modified phylogenetic trees with the online software Interactive tree of life (iTOL) [40].

### Protein domain diagram

The information about protein domains was collected from Pfam (http://pfam.xfam.org/), InterPro (http://www.ebi.ac.uk/interpro/) and UniProt (https://www.uniprot.org/). The illustration was depicted with IBS [41].

### RNA-seq data of Arabidopsis various tissues

The RNA-seq data of the Arabidopsis *RNHLS* genes were downloaded from TraVA (Transcriptome Variation Analysis, http://travadb.org/), except for which were not available [42]. The RNA-seq of rice *RNHLS* genes was downloaded from the Rice Genome Annotation Project (http://rice.plantbiology.msu.edu/expression.shtml). The heatmap was visualized using Morpheus online software with default settings. GO analysis of highly expressed genes in meristems was performed on the online database agriGO v2.0 [43].

## Results

### Species selected to cover the characters of RNHLS proteins in plants

To determine the representative plant species for the analysis of RNHLS proteins, we investigated co-occurring domain families with the value of the RNHLS proteins co-occurred with other protein domain families in 33 plant species (Fig. 1A, and Fig. S1) [29]. The result showed that most of the plant species had the co-occurrence of the RNHLS domain with a DNA polymerase domain, a nuclease, and nucleotide triphosphate hydrolase domain (Fig. S1). We thus selected ten representative plant species according to the Pearson correlation scores and included the other three species, *C. sinensis*, *M. polymorpha*, and *K. flaccidum* to better represent the evolutionary tree of plants for further investigation (Fig. 1B) [44].

**Figure 1.**
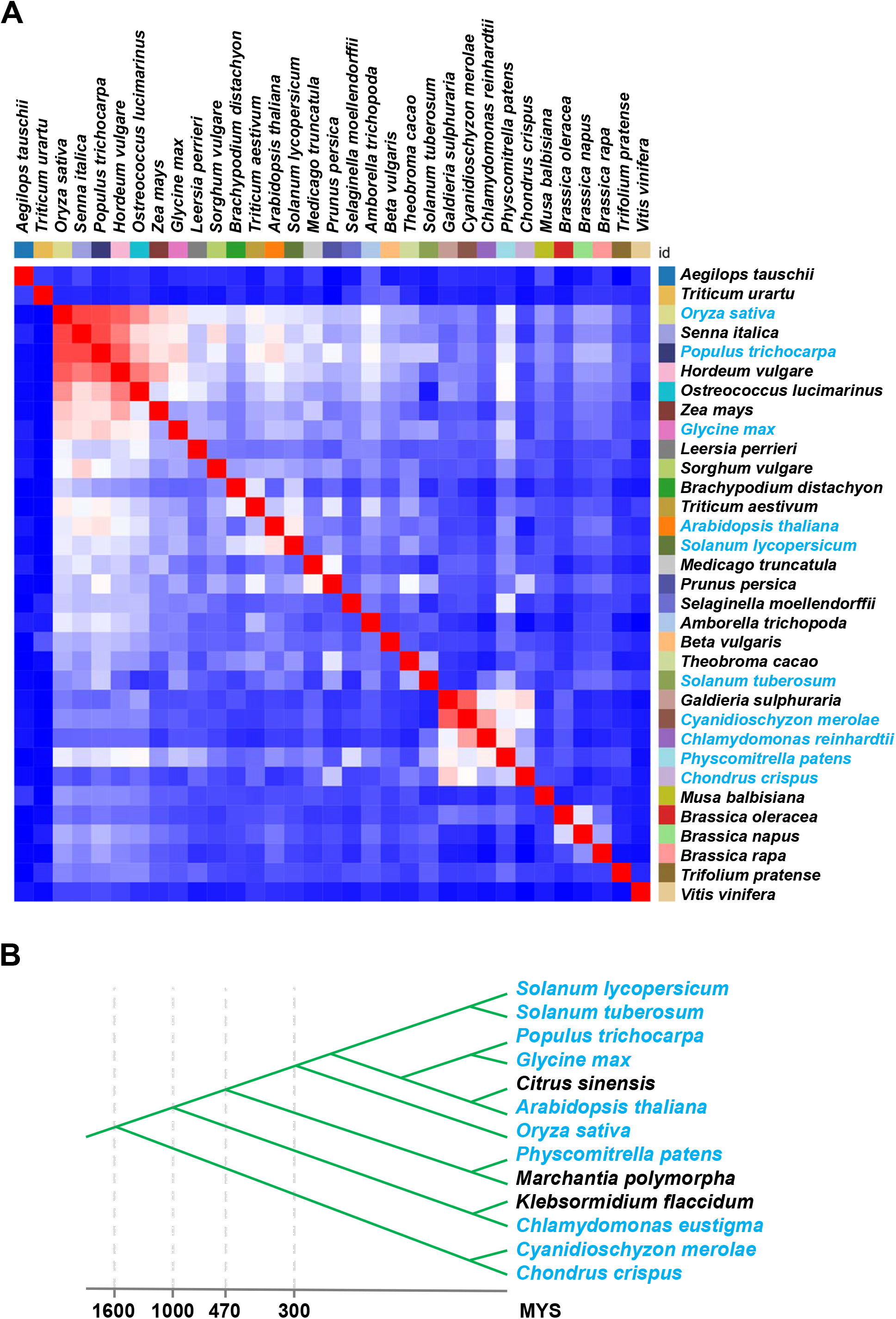
Selection of candidate species for investigation of plant RNHLS proteins. (A) The similarity of Gene3D domain family co-occurrence of RNHLS domain with other protein domains in all available plant species. The species to be investigated in this study were marked in blue font. The original matrix of different protein domains that co-occurred with the RNHLS domain was downloaded from Gene3D (http://gene3d.biochem.ucl.ac.uk). The similarity heatmap was performed in Morpheus (https://software.broadinstitute.org/morpheus/) using the default setting. (B) The cladogram of species for investigation of RNHLS proteins. The blue font is marked the selected species from the similarity matrix, while the black font is marked the species which are supplemented for better covering the plant evolutionary.

### A global overview of the RNHLS protein clusters

To understand the function and distribution of the RNHLS proteins in the 13 plant species, we acquired 6193 proteins from the protein database InterPro [45]. Next, we conducted a clustering analysis with protein sequence similarity to elucidate the relationship among them. We first chose the representative proteins using CD-HIT with a threshold of 60 percent and obtained 2066 representative RNHLS proteins. We then compared the similarity of these proteins with all RNHLS proteins using BLAST with an average pairwise E-value at least of 1×10^−18^. We found that they could be separated into 27 clusters in the network (Fig. 2). The majority of them might be Ty1/Copia or Ty3/Gypsy reverse transcriptases and other transposases or integrases. Besides, the second most significant part belongs to different types of nucleic acid endonucleases and exonucleases (Fig. 2). Additionally, the third part is Argonaute proteins and DNA polymerases. Therefore, the classification of these proteins was consistent with their biological function and confirmed that our SSN map could show the relevant information of the investigated RNHLS proteins. To further explore the relationship of the RNHLS proteins and the 27 subclusters, we constructed phylogenetic trees of total RNHLS proteins and each of the 27 subclusters. Consistent with the SSN network, the RNHLS proteins are diversified, and most of them are transposon polyprotein (Fig. S3 and Fig. S4).

**Figure 2.**
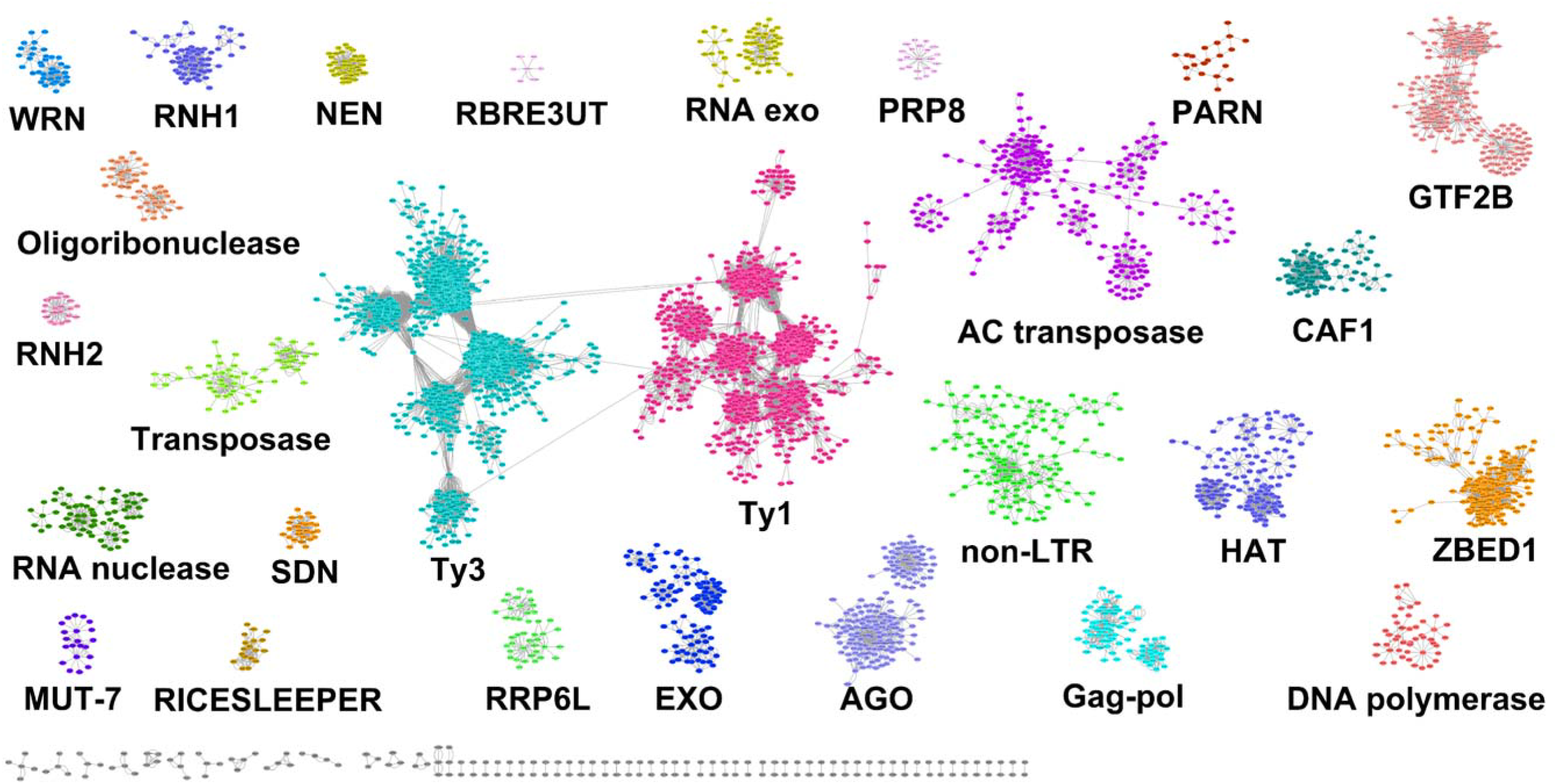
A representative SSN of the RNHLS proteins. A total of 4,755 protein sequences were depicted by 2,066 nodes (circles), which represent proteins sharing >50% sequence identity. Edges between nodes indicate an average pairwise BLAST E-value of at least 1 × 10^−18^. Node coloring represents subgroup classification. Names indicate subgroups that contain at least one protein with literature-documented functional annotations except for nodes marked with gray. The network was visualized by Cytoscape3.7.0 using the organic layout algorithm. WRN, Werner syndrome ATP-dependent helicase; SDN, small RNA degrading nuclease; RNH2, ribonuclease H2; AGO, Argonaute; RNH1, ribonuclease H1; RRP6L, RRP6-like protein; RBRE3UT, RBR-type E3 ubiquitin transferase; Exo, exonuclease; Ty1, Ty1/copia-element polyprotein; Ty3, Gypsy/Ty3 element polyprotein; non-LTR, putative non-LTR retroelement reverse transcriptase; CAF1, putative CCR4-associated factor 1 homolog; HAT, HAT dimerization domain-containing protein; GTF2B, general transcription factor 2-related zinc finger protein; ZBED1, zinc finger BED domain-containing protein DAYSLEEPER (Transposase-like protein DAYSLEEPER); Gag-pol, putative gag-pol polyprotein; PRP8, Pre-mRNA-processing-splicing factor 8.

Next, to further investigate the composition of these clusters at a species level, we indicated species information with different colors at the SSN map (Fig. S2). The results showed that most of the proteins from *O. sativa* are clustered in the Ty1/Copia and Ty3/Gypsy polyprotein clusters. Besides, the RNHLS proteins from *A. thaliana* and *O. sativa* could be gathered in different subclusters in the Ty1/Copia polyprotein group (Fig. S2). Moreover, proteins from *C. sinensis*, *C. crispus*, *S. lycopersicum,* and *eustigma* could also form subclusters in the Ty1/Copia polyprotein group. Interestingly, transposases from *C. crispus* and *M. polymorpha* could be separated into two relatively isolated groups (Fig. S2). Consistent with that, most of the transposon-related polyprotein clusters prefer to group in a species-dependent manner. Nonetheless, other clusters such as AGO, PRP8, DNA polymerase, and RNH2 might be distributed more evenly compared with transposon-related polyprotein clusters (Fig. S2). Furthermore, the AGO cluster could be divided into two sub-clusters consistent with their biological function (Fig. 2). Altogether, these results indicate that RNHLS proteins might be very diversified, and the evolution of transposon within each species might be exclusive.

### Phylogenetic tree of the Arabidopsis RNHLS proteins

To better understand the RNHLS proteins in plants, we used the RNHLS proteins from Arabidopsis to conduct an alignment and construct a phylogenetic tree (Fig. 3). Consistent with the protein clusters, the retrotransposon reverse transcriptases occupy a big part of the clades (Fig. 3). Additionally, there could be five clades of Ty1/copia reverse transcriptases, which is consistent with the cluster results that showed more than eight sub-clusters (Fig. 3). Likewise, the non-LTR retrotransposon reverse transcriptases comprise another three clades of the tree. On the contrary, the HAT transposon proteins might be clustered together in the tree (Fig. 3). Moreover, the CAF1 proteins are separated into two clades compared with MENs, SDNs, or exonucleases, which show only one clade on the tree (Fig. 3). Remarkably, the AGO family proteins could be separated into four clades of them with AGO1/5/10, AGO2/3, AGO4/6/8/9, and AGO7, which is identical to previous reports [46]. Nevertheless, various DNA polymerases with the RNHLS domain and the PRP8 family proteins are clustered into one clade (Fig. 3). Moreover, we found one type of RNHLS proteins from Ring-type E3 ubiquitin transferases (Fig. 3). Collectively, these results give us a comprehensive overview of the RNHLS proteins and the relationship among them in Arabidopsis.

**Figure 3.**
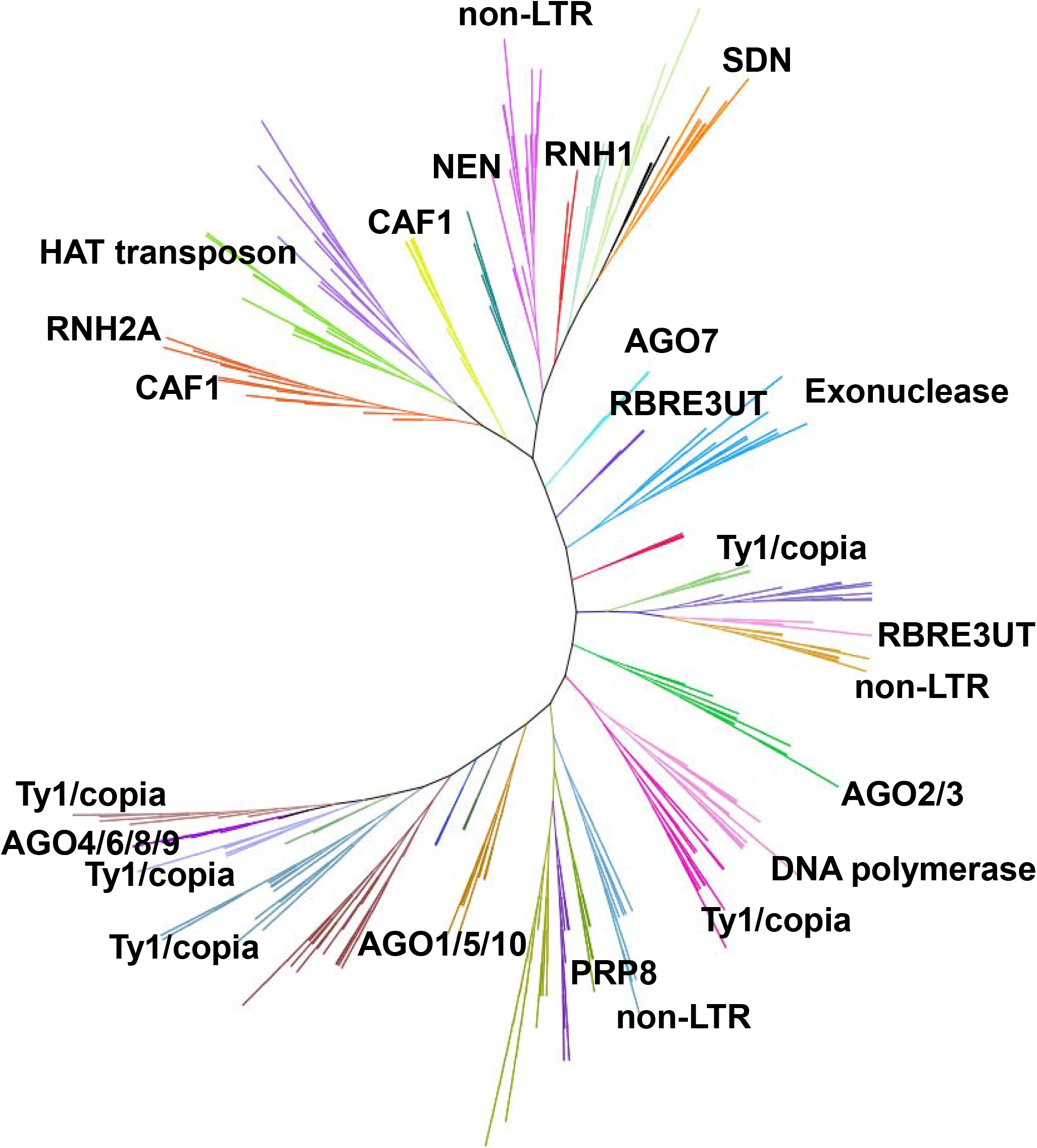
Phylogenetic tree of the Arabidopsis RNHLS proteins. A total of 691 Arabidopsis RNHLS protein sequences were aligned using Clustal Omega (https://www.ebi.ac.uk/Tools/msa/clustalo/). The phylogenetic tree was constructed using the results from this alignment and modified using iTOL (https://itol.embl.de/). Different types of nucleases and polymerases are clearly separated into distinct clades. The distribution of transposon-related proteins is more diversified than AGOs, SDNs, and MENs.

### The domain construction of different types of RNHLS proteins

To better characterize the properties of the RNHLS proteins, we chose the representative Arabidopsis RNHLS proteins in each of the clades to draw a protein domain diagram to indicate their domain composition (Fig. 4). The RNH1 contains three crucial domains, HBD (hybrid binding domain), Dis (disorder protein domain), and RNH domain (ribonuclease H domain). HBD is vital for RNA:DNA hybrid binding, while the RNH domain is for the catalytic activity [47]. The Dis domain provides the ability for RNH to catalyze a variety of hybrids without unloading (Fig. 4) [47]. Therefore, most RNHLS proteins could combine RNH domain with other domains to exert their function to overcome RNA:DNA hybrid during different biological processes. In addition, to analyze the relationship of the RNHLS domain in different RNHLS proteins, we performed a multiple sequence alignment with the truncated sequences of representative RNHLS proteins from Arabidopsis (Fig. 5). The results indicate that the conserved alpha-helix and beta-sheet domain present in these RNHLS proteins consistent with their annotated function in InterPro (Fig. 4 and Fig. 5).

**Figure 4.**
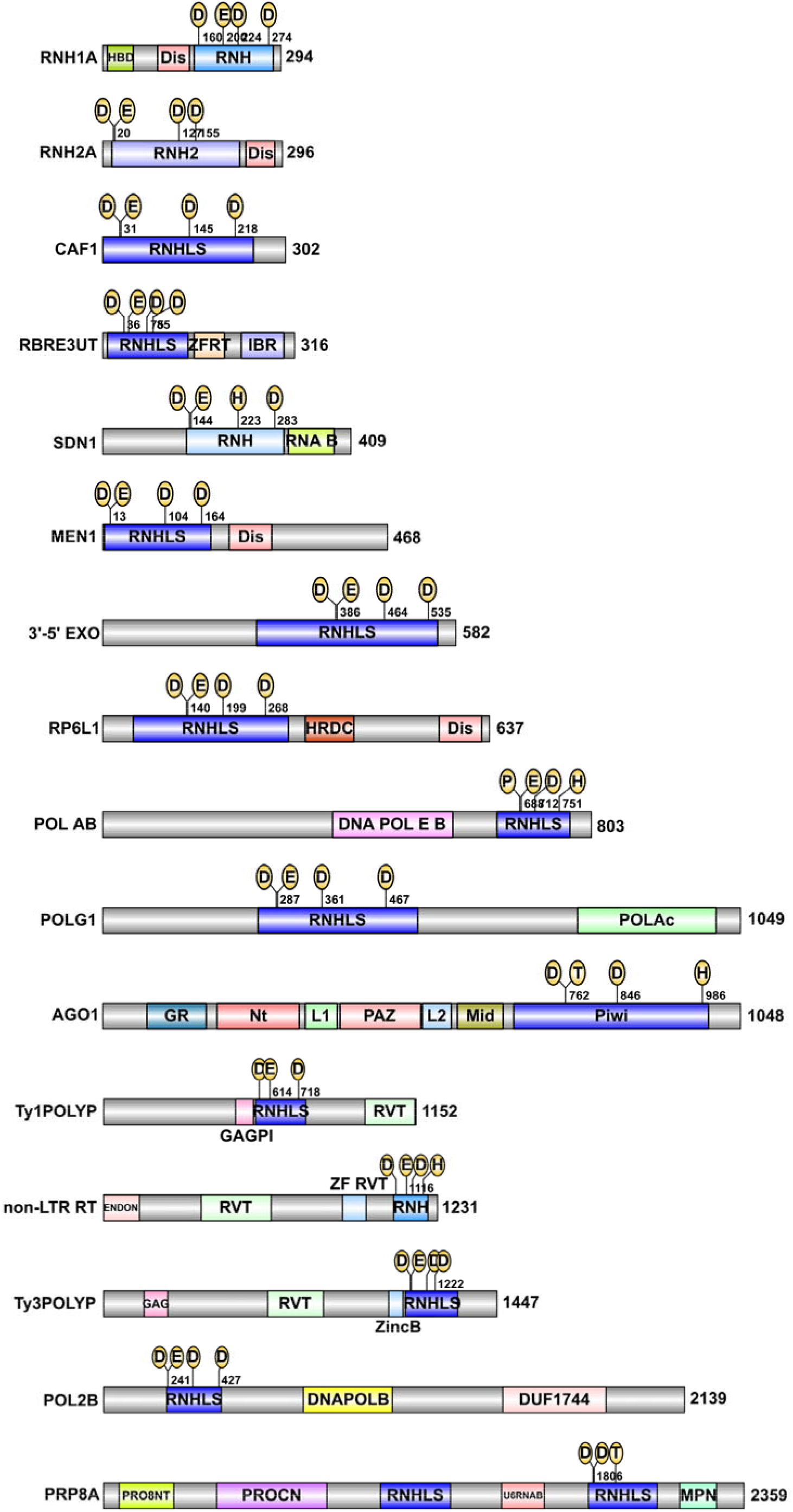
Protein domain diagram of Arabidopsis representative RNHLS proteins. The domain annotations of these proteins were combined from Pfam (http://pfam.xfam.org/), InterPro (http://www.ebi.ac.uk/interpro/) and UniProt (https://www.uniprot.org/). HBD, hybrid binding domain; Dis, disorder protein domain; RNH, ribonuclease H domain; RNHS, Ribonuclease H-like superfamily domain; ZFRT, zinc finger ring-type domain; IBR, In Between Ring fingers domain; RNA B, RNA binding domain; HRDC, helicase and RNase D C-terminal; DNA POL E/B domain, DNA polymerase alpha/epsilon subunit B domain; POLAc, DNA polymerase A domain; GR, glycine-rich domain; Nt, N-terminal domain; L1, linker 1 domain; AGO1, Argonaute 1, consists of GR, glycine-rich domain, Nt, N-terminal domain, L1, linker 1 domain; PAZ, PAZ domain; L2, linker 2 domain; Mid, Mid domain; Piwi, Mid domain; Ty1POLYP, Ty1/copia-element polyprotein; GAGPI, GAG-pre-integrase domain; RVT, reverse transcriptase, RNA-dependent DNA polymerase domain; ENDON, Endonuclease superfamily domain; ZF RVT, reverse transcriptase zinc-binding domain; Ty3POLYP, Ty3/Gypsy-element polyprotein; GAG, retrotransposon gag domain; ZincB, integrase zinc-binding domain; POL2B, DNA polymerase epsilon catalytic subunit; DNAPOLB, DNA-directed DNA polymerase family B multifunctional domain; DUF1744, domain of unknown function 1744 domain; PRP8, Pre-mRNA-processing-splicing factor 8; PRO8NT, N terminus of PRP8 domain; PROCN, central domain of PRP8 domain; U6RNAB, U6-snRNA-binding of PRP8 domain; MPN (Mpr1, Pad1 N-terminal) domain. D, aspartic acid. E, glutamic acid. H, histidine. T, threonine. The numbers beside the protein indicate the amino acid number of that protein.

**Figure 5.**
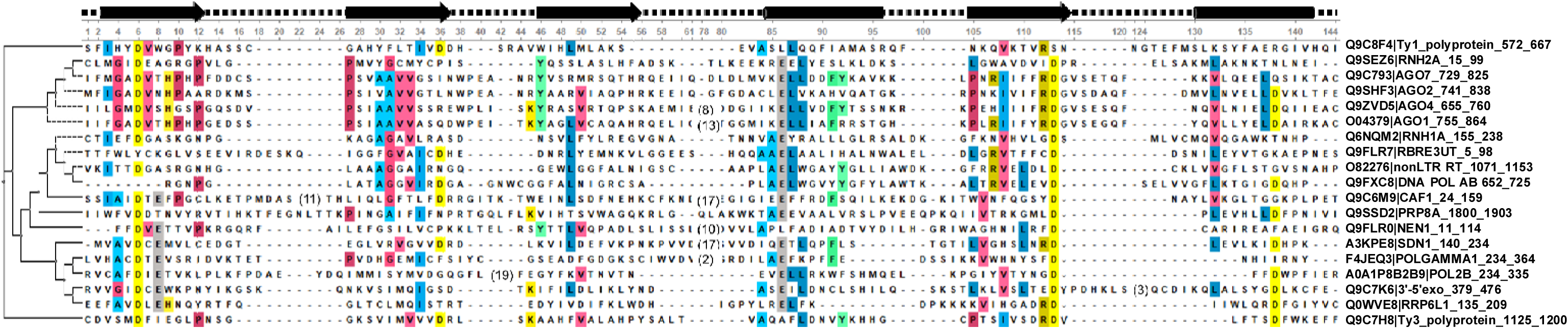
Multiple sequence alignment of Arabidopsis representative RNHLS proteins. The RNHLS domains of each protein were selected for this alignment. Each of RNHLS domains was predicted the secondary structure using Jpred4 (http://www.compbio.dundee.ac.uk/jpred/index.html). The alignment was visualized using UGENE and adjusted manually.

### RNHLS proteins are abundant in plant meristems

To predict the biological function of these RNHLS proteins in plants, we first analyzed the expression pattern of the RNHLS proteins in Arabidopsis and rice using RNA-seq data (Fig. 6 and Fig. S5). In Arabidopsis and rice, various *RNHLS* genes were expressed in different tissues and could be grouped. Moreover, we found these genes in Arabidopsis are mainly expressed at meristem tissues, such as shoots, root meristems, and inflorescences (Fig. 6A). Additionally, some of them are mainly expressed in anthers and pistils (Fig. 6A). Nonetheless, different types of *RNHLS* genes show distinct expression patterns in the young and mature anthers (Fig. 6A). Meanwhile, the dry seeds accumulate another group expression of *RNHLS* genes (Fig. 6A). However, some genes could be expressed in senescent tissues probably due to programmed cell death accompanied by nucleic acid degradation (Fig. 6A). Besides, we performed a GO analyzer of the meristem expressed genes which showed that they are mainly involved in nucleic acid metabolisms (Fig. 6B). Furthermore, post-emergence and pre-emergence inflorescences are grouped with the different expressions of *RNHLS* genes in rice similar to that in Arabidopsis (Fig. S5). Intriguingly, there is another group of *RNHLS* expressed in the embryo-25 dap (days after plant) (Fig. S5). Moreover, half of the investigated *RNHLS* genes are expressed at the four-leaf stage (Fig. S5). Taken together, the *RNHLS* genes are expressed throughout the life cycle of Arabidopsis and rice, which is in accordance with their crucial biological function. These results collectively indicate the importance of *RNHLS* genes in plants.

**Figure 6.**
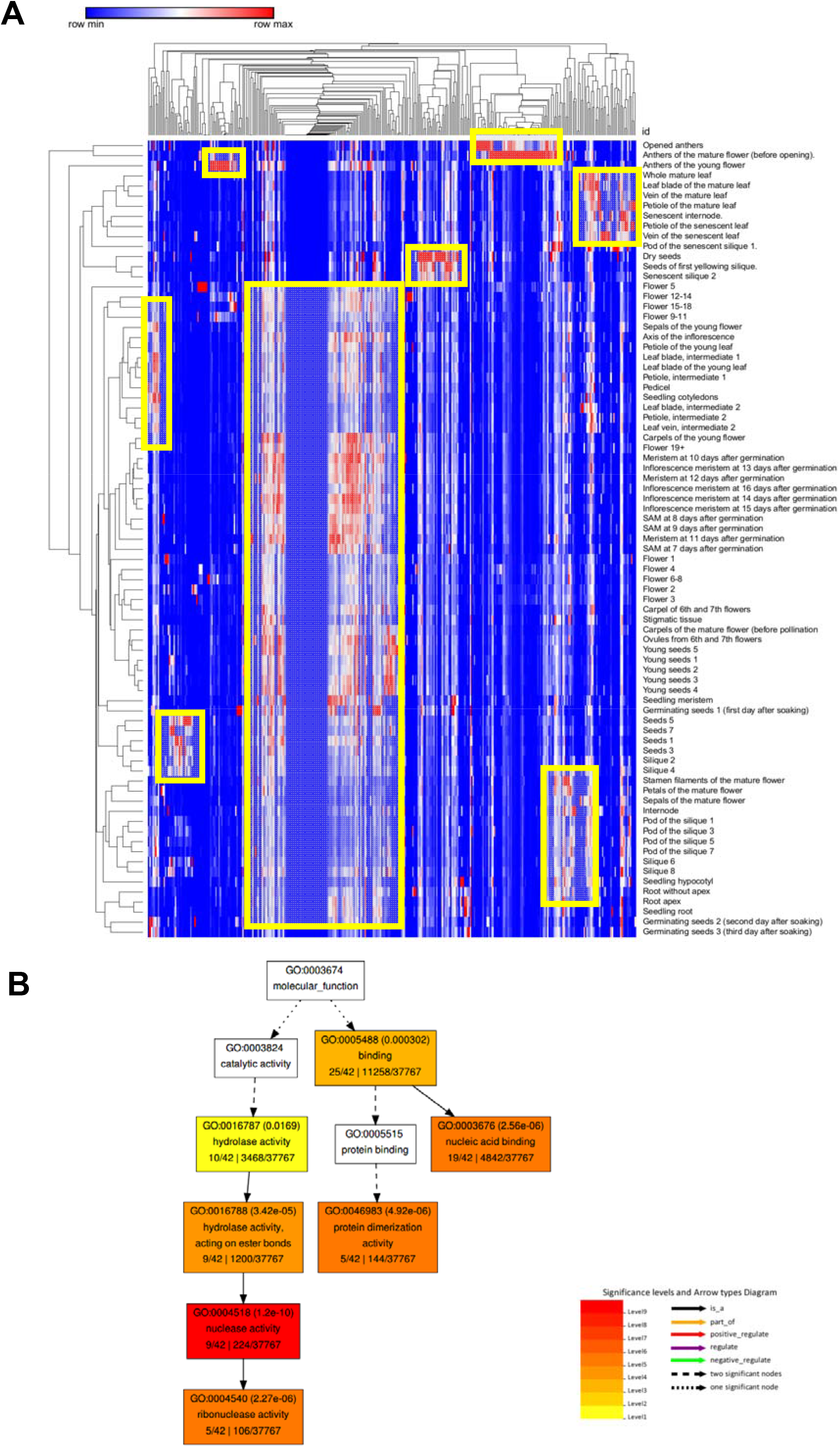
Tissue-specific expression of Arabidopsis *RNHLS* genes. (A) The transcriptome of *RNHLS* genes was downloaded from TRAVA (http://travadb.org/browse/) except for some of the genes that could not be retrieved from that database. These genes are mainly expressed at the meristems such as flower inflorescences and root meristems. The yellow rectangles indicate the clusters of genes highly expressed in these tissues. (B) GO annotation of the genes expressed in Arabidopsis meristems. These genes are mainly involved in nucleic acid metabolism.

## Discussion

In this study, we characterized 6193 RNHLS proteins in 13 plants. Representative 2066 RNHLS proteins are clustered into 27 clusters, including reverse transcriptases, transposases, integrases, nucleic acid endonucleases, exonucleases, key spliceosome components Prp8, as well as other RNA nucleases. Moreover, we showed that the RNHLS domain is combined with other protein domains in different RNHLS proteins. Besides, the catalytic alpha-helix and beta-sheet domain are conserved among them in Arabidopsis. Particularly, *RNHLS* genes are actively expressed in meristems, flower tissues, and other metabolic tissues. Various *RNHLS* genes expressed in different tissues could be grouped.

The selected 13 plant species represent the evolutionary process of plants (Fig. 1B) [44]. The distribution of RNHLS proteins in all species of plants suggests they might be one of the evolutionarily oldest proteins [16]. Among the 27 clusters of the RNHLS proteins, DNA polymerases, GTF2Bs, CAF1s, PRP8s, RRP6Ls, AGOs, and EXOs perform their function in basic nucleic acid metabolic processes which is conserved in the plant kingdom [15]. A previous study showed that WRNs can digest the RNA moiety of the RNA:DNA hybrid [48]. Recently, WRN is found to be involved in R-loop-related genome stability [49]. Moreover, SDN proteins could trim dsRNA and be involved in degrading small RNAs thus raising the possibility of trimming RNA strands in RNA:DNA hybrids as RNA-DNA hybrids might be high structural similar to dsRNAs [50]. Interestingly, the family number of TE-related RNHLS proteins might be more extensive, especially for Ty1/Copia and Ty3/Gypsy reverse transcriptases (Fig.2 and Fig. S4). However, some of the TE-related RNHLS proteins are not present in all selected plants, such as RICESLEEPERs, which only present in rice (Fig. S2). Moreover, Ty1/Copia, Ty3/Gypsy, and Non-LTR reverse transcriptases are mainly enriched in rice, Arabidopsis and other higher plants (Fig. 2 and Fig. S2). That might be in line with the size and composition of their genomes [44]. As retrotransposons are abundant in genomes of higher plants, the TE-related RNHLS proteins may play major roles in driving the genetic diversity and evolution of these organisms [13, 51].

The RNHLS family might be a large group of evolutionarily related but strongly diverged protein families [15]. The first RNHLS protein identified is RNase H from *Escherichia coli* [52]. There are two types of RNase H enzymes, which differ in structure and substrate specificity. RNase H1 is monomeric, whereas RNase H2 is monomeric in bacteria but composed of three subunits in eukaryotes: RNH2A (the catalytic subunit), RNH2B and RNH2C. Both types of RNase H enzymes can remove RNA-DNA hybrids in addition to having different specialized roles. Surprisingly, the phylogenetic analysis of the RNHLS proteins in Arabidopsis showed that some proteins with similar functions were clustered into different groups according to their sequence, such as AGO proteins. Besides, some proteins from different protein families were clustered together, such as some Ty1/Copia and non-LTR reverse transcriptase proteins. All these results indicated that both amino acid constructions and motif compositions of RNHLS proteins are diversified. In addition, domain analysis of the representative proteins showed that all the RNHLS proteins might have RNH domains except AGOs, which uses a PIWI domain instead (Fig.4). The Piwi domain has been reported to adopt an RNase H-like fold, which enables some, but not all, AGO proteins to cleave target RNAs complementary to their bound sRNAs [19]. Furthermore, a catalytic triad (Asp-Asp-His/Asp, DDH/D) is generally thought to be responsible for slicer activity which presents in both RNH domains and PIWI domains [53]. Thus, further studies are needed to assess whether AGOs are capable of destroying RNA strands in RNA-DNA hybrids or as a sensor for RNA-DNA hybrids. In addition, the RNHLS proteins are abundant in meristems, flower tissues, and other metabolic tissues while various *RNHLS* genes expressed in different tissues could be grouped which indicates that they might be critical for the development of plants (Fig. 6). As RNHLS proteins work on nucleic acid metabolism especially on R-loop homeostasis, this study characterized the RNHLS proteins in plants and provide new direction on R-loop regulation and function identification study in the future.

## Conclusions

In this study, we employed the bioinformatics approaches to analyze the RNHLS proteins in 13 representative plant species. An SSN network showed that these proteins can be separated into 27 clusters and consistent with their biological functions. Moreover, the *RNHLS* genes expression pattern during Arabidopsis and rice development further revealed that these proteins are mainly expressed in meristems and metabolic tissues. These results shed light on our understanding of R-loop regulation in plants.

## Supplementary material

**Supplementary material 1:** Supplementary figure 1 to supplementary figure 5. (PDF 2560 kb)

**Supplementary material 2:** A representative SSN of the RNHLS proteins. (CYS 4687 kb)

**Supplementary material 3:** A representative SSN of the RNHLS proteins colored with species information. (CYS 4657 kb)

**Supplementary material 4:** Supplementary table 1. Gene3D domain family matrix of RNHLS domain that co-occurs with other protein domains. (XLSX 52.4 kb)

**Supplementary material 5:** Supplementary table 2. RNHLS protein information used in this study. (XLS 668 kb)

**Supplementary material 6:** Supplementary table 3. tissue-specific expression data of *RNHLS* genes of Arabidopsis and rice. (XLSX 187 kb)

## Abbreviation

AGO: Argonaute
CAF1: putative CCR4-associated factor 1 homolog
Dis: disorder protein domain
DUF1744: domain of unknown function 1744 domain
Exo: exonuclease
Gag-pol: putative gag-pol polyprotein
GTF2B: general transcription factor 2-related zinc finger protein
HBD: hybrid binding domain
non-LTR: putative non-LTR retroelement reverse transcriptase
Prp8: pre-mRNA-processing-splicing factor 8
RBRE3UT: RBR-type E3 ubiquitin transferase
RNase H: ribonucleases H
RNH2: ribonuclease H2
RNHLS: RNase H-like superfamily
RRP6L: RRP6-like protein
SDN: small RNA degrading nuclease
SSN: sequence similarity network
WRN: Werner syndrome ATP-dependent helicase
ZBED1: zinc finger BED domain-containing protein DAYSLEEPER (Transposase-like protein DAYSLEEPER).

## Author contributions

Q.S. conceptualized the study. S.L. analyzed the sequences and prepared the figures, and prepared the manuscript with K.L. and Q.S.

## Acknowledgments

This work was funded by grants from the Ministry of Science and Technology of China (grant no. 2016YFA0500800 to Q.S.); and the National Natural Science Foundation of China (grant nos. 91740105 and 31822028 to Q.S.). The Sun Lab was supported by Tsinghua-Peking Joint Center for Life Sciences, Tsinghua University Initiative Scientific Research Program, and the 1000 Young Talent Program of China. K.L. was supported by the postdoctoral fellowship from Tsinghua-Peking Joint Center for Life Sciences.

